# MRIQC Web-API: Crowdsourcing image quality metrics and expert quality ratings of structural and functional MRI

**DOI:** 10.1101/216671

**Authors:** O. Esteban, RW. Blair, DM. Nielson, JC. Varada, S. Marrett, AG. Thomas, RA. Poldrack, KJ. Gorgolewski

## Abstract

The MRIQC Web-API is a resource for scientists to train new automatic quality classifiers. The MRIQC Web-API has collected more than 30K sets of image quality measures automatically extracted from BOLD and T1-weighted scans using MRIQC. MRIQC is an automated MRI Quality Control tool, and here we present an extension to crowdsource these quality metrics along with anonymized metadata and manual quality ratings. This new resource will allow a better understanding of the normative values and distributions of these quality metrics, help determine the relationships between image quality and metadata such as acquisition parameters and finally, provide a cost-effective, easy way to annotate the quality of a large number of cross-site MR scans.

## Introduction

Neuroimaging is experiencing the data deluge that is occurring across the sciences. The numbers of MR images of the human brain that are daily acquired make manual quality control of those data impractical. MRIQC^1^ was proposed recently to advance the automated quality control (QC) of T1-weighted, T2-weighted and BOLD (blood-oxygen-level dependent) MRI. However, to ensure the generalizability in predicting quality based on the image quality metrics (IQMs) extracted with MRIQC, it is necessary to annotate a very large number of images. Here, we present a lightweight web-service tailored to MRIQC that collects those image quality metrics in a large MRI-quality database. Additionally, the MRIQC Web-API is capable of storing quality ratings made by experts with an addon to the visual HTML reports generated by MRIQC.

## Methods

The Web-API is an addon to any automated QC software that generates image quality metrics (IQMs). The IQMs are collected in a publicly accessible database, that can be queried at https://mriqc.nimh.nih.gov/. The overall framework involves the following workflow (see Figure 1):

1. <u>Execution of MRIQC and submission of IQMs:</u> T1w, T2w and BOLD images are processed with MRIQC that computes a number of IQMs (a broader description of these measures are available at the documentation site, http://mriqc.rtfd.io/en/stable/measures.html). The IQMs are collated in a JSON record, which MRIQC automatically submits to a Representational state transfer (REST) or RESTful endpoint of the Web-API. These JSON records are then stored, including a unique identifier generated from the actual data array of the original image and additional anonymized metadata and provenance. Users can opt-out if they do not wish to share their IQMs. MRIQC is an easy-to-use, multiplatform software that can be installed natively in Unix systems or run in MacOSX and Windows through Docker. MRIQC can also run in HPC clusters where Docker is not available through Singularity^2^. Finally, it can be directly executed without installation on datasets stored on OpenNeuro (http://openneuro.org)^3^
2. <u>Visualization of the individual reports:</u> MRIQC generates dynamic HTML (hypertext markup language) reports that speed up the visual assessment of each image of the dataset by the expert. MRIQC version 0.9.11 includes a widget (see Figure 2) that allows the researcher to assign a quality rating to the image under review.
3. <u>Crowdsourcing expert quality ratings:</u> the RESTful endpoint receives the quality ratings, which are linked to the original image via their unique identifier.

**Figure 1.**
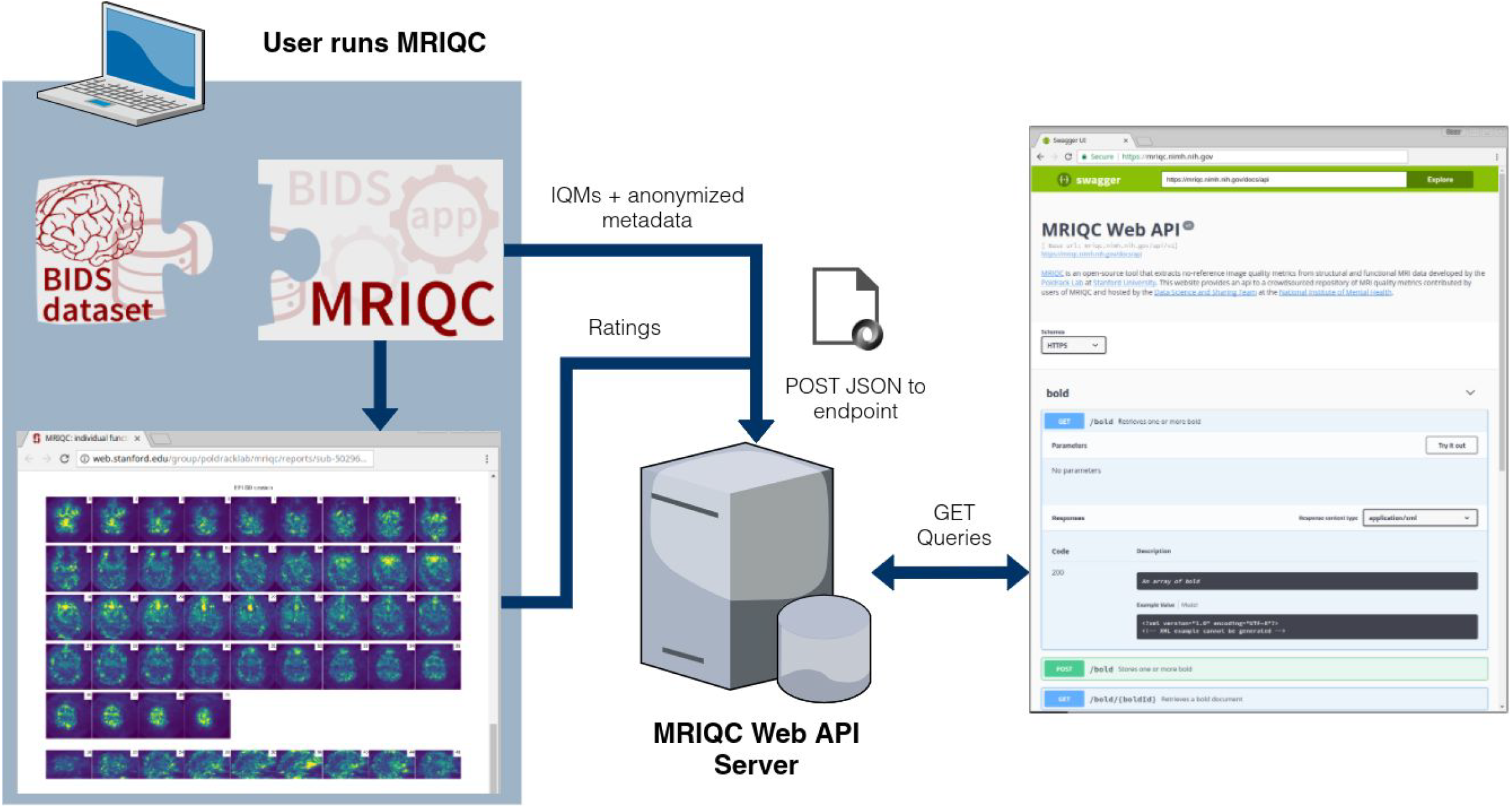
A BIDS dataset is processed with MRIQC. Processing finishes with a POST request to the MRIQC Web API endpoint with a payload containing the image quality metrics (IQMs) and some anonymized metadata (e.g. imaging parameters, the unique identifier for the image data, etc.) in JSON format. Once stored, the endpoint can be queried to fetch the crowdsourced IQMs. Finally, a widget (Figure 2) allows the user to annotate existing records in the MRIQC Web API.

**Figure 2.**
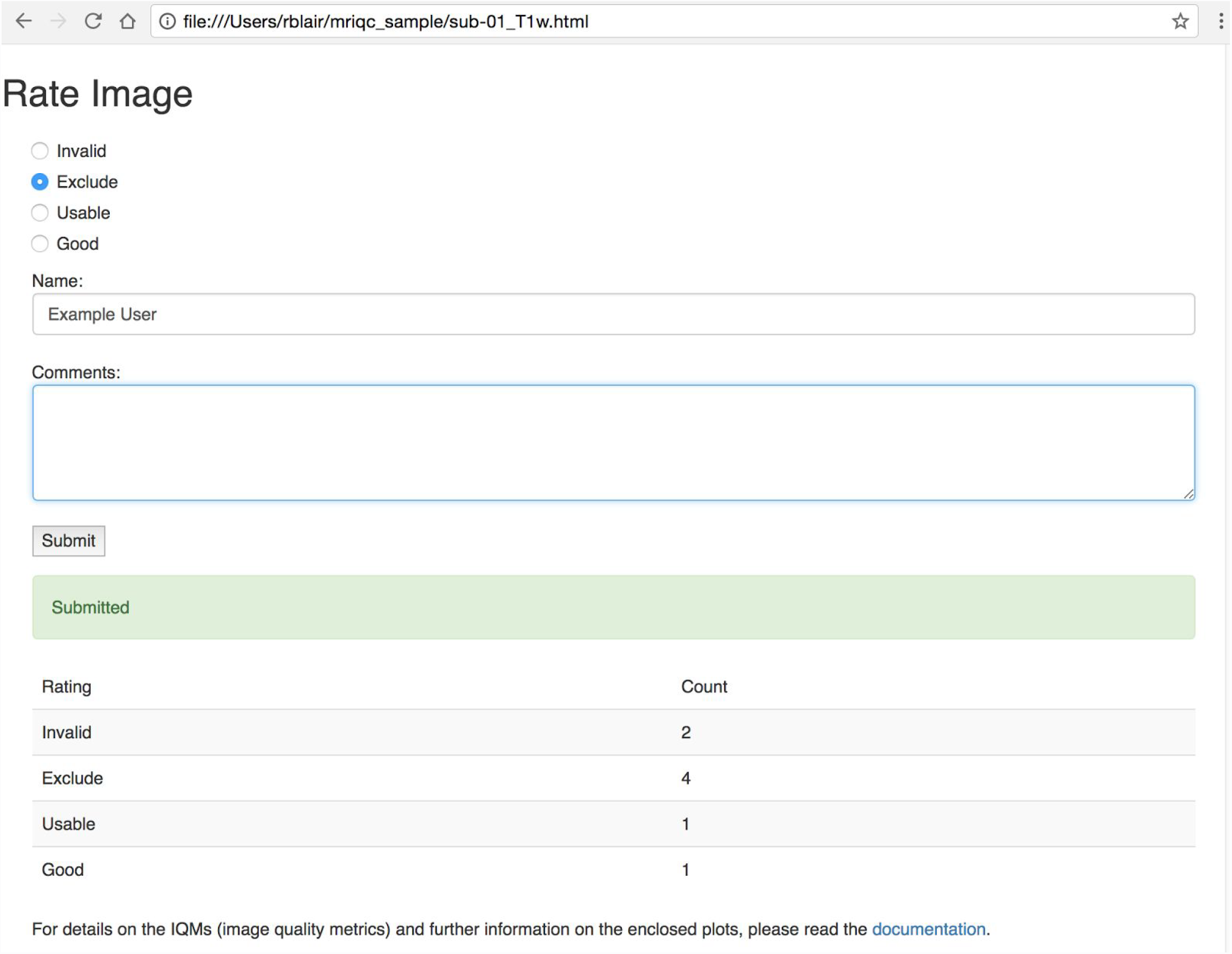
A new widget included by the individual image report allows users to annotate quality of existing records in the MRIQC Web API.

## Results

At the time of writing, the MRIQC Web-API database contains more than 15 thousand records from T1w images and 22 thousand from BOLD scans. Figure 3A shows the historical evolution, and Figure 3B presents the distributions of some IQMs extracted from T1w and BOLD images collected so far. An example of Jupyter Notebook to query the API is maintained in the GitHub repository^4^. Additionally, the crowdsourcing of expert quality ratings is fully functional.

**Figure 3.**
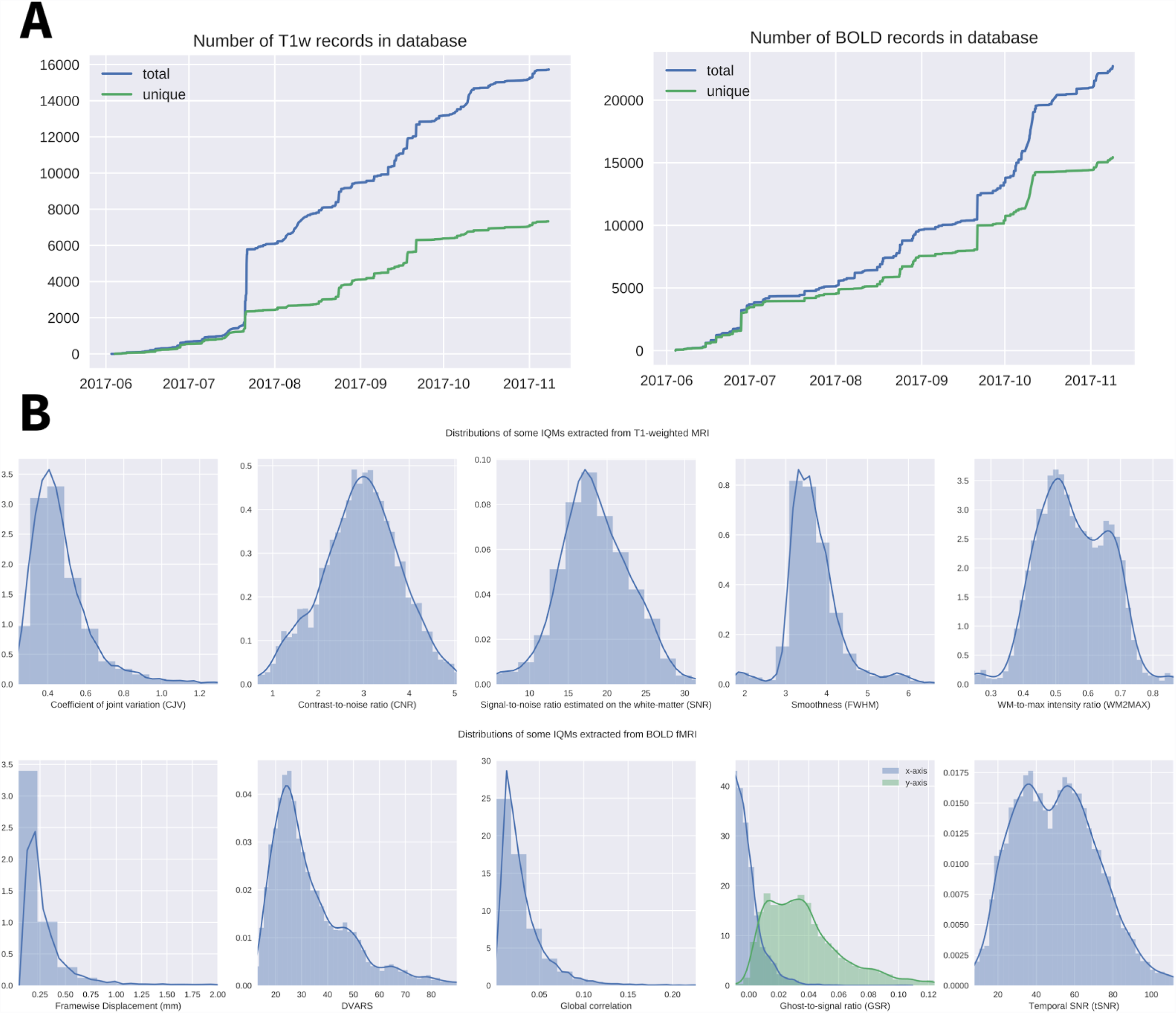
A) Cumulative number of records, by modality; B) Example of distributions of several image quality metrics (IQMs) for T1-weighted images (first row) and for BOLD fMRI (second row).

## Discussion and conclusion

The MRIQC Web-API is a resource for scientists to train new automatic quality classifiers. Features for automated learning frameworks are collected through a web service that stores image quality metrics (IQMs) extracted automatically by neuroimaging software (e.g. PCP-QAP^5^) from MRI scans. The web service is also able to permanently store linked quality ratings made by human experts after the visual inspection of the reports generated by MRIQC. However, the interpretation and most importantly, the normative values of these metrics are widely unknown and are highly dependent on the imaging center, hardware details such as vendor, and the particular scanning parameters. By crowdsourcing the IQMs we expect to advance the interpretation of these metrics. The cumulative addition of expert ratings to annotate the IQMs will improve the prediction accuracy of automatic quality classifiers. The process of individually rating each MR scan is tedious and time consuming since it involves the careful inspection of data and recording the assessment in some sort of database. MRIQC eases this process with thorough visual reports that speed up the assessment of images. The rating widget simplifies the final step of storing the manual quality rating. Integrating MRIQC in science-as-a-service platforms like OpenNeuro (http://openneuro.org) streamlines the overall work-flow and will allow for a more systematic and rigorous assessment of MR quality.

